# Complete Genome Sequences of Nine Cluster A Mycobacteriophages: AgentM, Ajay, Aragog, Archetta, ForGetIt, Koko, Ph8s, Phlorence, and Wilkins

**DOI:** 10.1101/2020.11.29.402826

**Authors:** Rachel M. Lipman, Amanda C. Freise, Andrew Kapinos, Canela Torres, Krisanavane Reddi, Jordan Moberg Parker

## Abstract

**Background:** Bacteriophages are ubiquitous, highly diverse, and relatively understudied. Growing interest in phage therapy has underscored the importance of isolating and characterizing novel bacteriophages. The Science Education Alliance-Phage Hunters Advancing Genomics and Evolutionary Science (SEA-PHAGES) program aims to address this need by involving undergraduates around the world in authentic research.

**Materials and Methods:** Nine novel mycobacteriophages - AgentM, Ajay, Aragog, Archetta, ForGetIt, Koko, Ph8s, Phlorence, and Wilkins - were isolated from soil samples in southern California using host *Mycobacterium smegmatis* mc2155. Each purified phage was characterized using transmission electron microscopy and genome sequencing and annotation.

**Results:** All nine bacteriophages were placed into mycobacteriophage Cluster A based on nucleotide similarity with other phages. The average genome length of all nine phages was 50,706 bp. On average, each phage had 87 total coding genes and GC content between 60-63%, consistent with that of other Cluster A phages. Transmission electron microscopy of phage particles revealed they had icosahedral heads and long, flexible tails, consistent with that of the *Siphoviridae* family. Plaque morphology and genome analysis confirms the nine novel phages are temperate as expected of Cluster A phages.

**Conclusions:** AgentM, Ajay, Aragog, Archetta, ForGetIt, Koko, Ph8s, Phlorence, and Wilkins are all mycobacteriophages that belong to the *Siphoviridae* family. Comparative genomic analyses revealed genetic mosaicism and diversity among these Cluster A phages. The discovery of these novel phages expands on the existing library of mycobacteriophage genomes.

## Introduction

The Howard Hughes Medical Institute Science Education Alliance-Phage Hunters Advancing Genomics and Evolutionary Science (SEA-PHAGES) initiative is an international research program that harnesses the power of undergraduate researchers to isolate, sequence, and characterize actinobacteriophages across the world for potential real-world applications [1]. To date, students in the program have isolated over 18,600 actinobacteriophages and sequenced over 3,500 (https://PhagesDB.org). These phages have already begun to have translational impact; several novel bacteriophages previously discovered by students involved in the SEA-PHAGES program were recently used to treat a disseminated drug-resistant *Mycobacterium abscessus* infection in a cystic fibrosis patient [2].

Here we report the isolation and characterization of nine novel Cluster A mycobacteriophages – AgentM, Ajay, Aragog, Archetta, ForGetIt, Koko, Ph8s, Phlorence, and Wilkins – by students at the University of California, Los Angeles in collaboration with SEA-PHAGES. These bacteriophages were isolated using bacterial host *Mycobacterium smegmatis* mc2155, a nonpathogenic and easily experimentally manipulated species in the *Mycobacterium* genus.

## Materials and methods

### Phage collection, isolation, and purification

All soil samples except that which produced phage Ajay were collected from the greater Los Angeles, CA area. Phage Ajay was isolated from a sample obtained 40 miles further east in Eastvale, CA. Phage extraction occurred through either direct or enriched isolation, as detailed in the SEA-PHAGES Discovery Guide https://seaphagesphagediscoveryguide.helpdocsonline.com/).

Phage Ajay was isolated via direct isolation by combining soil and 10X 7H9 Middlebrook enrichment broth (supplemented with albumin, dextrose, carbenicillin [50 μg/ml] and cycloheximide [10 μg/ml]) in a 15 mL conical tube and shaking at 250 rpm for 1-2 hours. After incubation, the mixture was filter sterilized. 500 μL of the mixture was added to 250 μL of host bacterium *M. smegmatis* mc^2^155 and incubated for 5-10 minutes. Lastly, 3 mL of molten top agar was added to the inoculated host tube, mixed, and then poured onto an agar plate. The plates were incubated at 30°C and observed for plaques 24-48 hours later. A turbid plaque was picked and purified via two consecutive plaque assays to produce a single phage population. Purified lysates were amplified via additional plaque assays to produce a high-titer phage lysate.

The other eight phages described were discovered through enriched isolation. Soil samples were similarly incubated with enrichment broth and a 48-hour culture of *M. smegmatis* and shaken for 2 days. After incubation, the sample mixture was centrifuged at 2,000 x g for 10 minutes. Supernatant was removed and filter sterilized. 10 μL of each supernatant sample was spotted on to a lawn of *M. smegmatis*. Plates were incubated at 30 or 35°C and plaques were picked and purified as above.

### Transmission electron microscopy

Phage lysates were pipetted onto a carbon-coated Formvar grid. Grids were then stained with 1% uranyl acetate (UA), dried, and viewed using a FEI T12 electron microscope (Thermo-Fisher, USA). Each phage was observed at multiple magnifications.

### DNA extraction and genome sequencing

Viral DNA was extracted and purified using the Promega Wizard DNA Clean-Up Kit (Product #A7280). Whole genome sequencing was conducted at the North Carolina State Genomic Sciences Laboratory, the Pittsburgh Bacteriophage Institute, or the UCLA Genotyping and Sequencing Core, using either Illumina sequencing or 454 GS FLX pyrosequencing. The sequencing libraries were created using the NEBNext® Ultra™ II DNA Library Prep Kit (New England Biolabs, MA, USA). Coverage values are reported in Table 1. Genome assembly was performed as described previously [3].

**Table 1.** Novel Cluster A mycobacteriophage sequencing information

| Phage Name | Cluster | GenBank Accession no. | Sequencing Coverage (x) | 3' Sticky Overhang Sequence |
| --- | --- | --- | --- | --- |
| Ajay | A1 | MN062710 | 4,218<sup>a</sup> | CGGATGGTAA |
| Wilkins | A1 | MG099951 | 32<sup>c</sup> | CGGATGGTAA |
| Ph8s | A2 | MG099947 | 14,824<sup>b</sup> | CGGTCGGTTA |
| AgentM | A5 | MG099934 | 62<sup>c</sup> | CGGGAGGTAA |
| Aragog | A5 | MG099937 | 76<sup>c</sup> | CGGGAGGTAA |
| Archetta | A5 | MG099938 | 103<sup>c</sup> | CGGGAGGTAA |
| ForGetIt | A5 | MG099944 | 144<sup>c</sup> | CGGGAGGTAA |
| Phlorence | A5 | MG099949 | 92<sup>c</sup> | CGGGAGGTAA |
| Koko | A6 | MG099945 | 11,617<sup>b</sup> | CGGTCGGTTA |

<sup>a</sup> Sequencing performed at the NC State Genomic Sciences Laboratory (Illumina)
<sup>b</sup> Sequencing performed at the Pittsburgh Bacteriophage Institute (Illumina)
<sup>c</sup> Sequencing performed at the UCLA Genotyping and Sequencing Core (454 GS FLX pyrosequencing)

### Genome Annotation

All phage genomes underwent auto and manual annotation. DNA Master (http://cobamide2.bio.pitt.edu/computer.htm) was used for auto-annotation. Glimmer [4] and GeneMark [5] were used to predict coding potential and identify open reading frames. ARAGORN [6] and tRNAScan-SE [7] were used to predict and trim tRNAs for each phage. Manual annotation was performed using PECAAN (http://pecaan.kbrinsgd.org), Phamerator [8] and Starterator [9]. NCBI BLAST [10], HHPred [11], and the Conserved Domain Database [12] were used to predict gene functions. TMHMM [13] was used to predict transmembrane domains.

## Results

TEM imaging of each phage revealed that they all belonged to the *Siphoviridae* family, based on their long, flexible tails (Figure 1).

**Figure 1.**
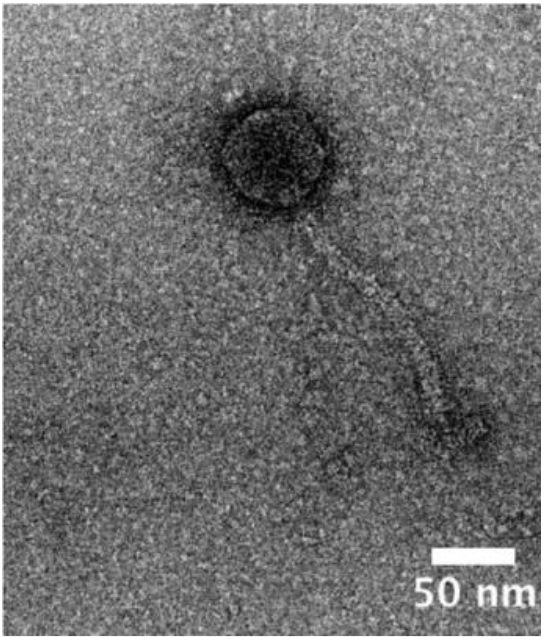
Isolated Cluster A mycobacteriophages have *Siphoviridae* morphology. Representative image of Cluster A phage Wilkins at 26,000X magnification, showing its long flexible tail and icosahedral head. All phages described had similar features.

The nine novel mycobacteriophages were placed into Cluster A based on their nucleotide similarity with other phages. Ajay and Wilkins were placed into subcluster A1, Ph8s into A2, Agent M, Aragog, Archetta, ForGetIt, and Phlorence into A5, and Koko into A6. All nine phages share a 3’ sticky end overhang of 10 base pairs, and phages in the same subcluster have the same overhang sequence (Table 1). The average genome size was 50,706 bp. The nine phages had a median of 84 total protein coding genes (range: 76-102), 0-3 tRNAs, and an average GC content of 61.7% (range: 60.6-63.9%) (Table 2). Each of these values is consistent with overall Cluster A genomes (https://phagesdb.org/clusters/A/). In all nine phages, structural and assembly genes were conserved on the left arm of the genome [14] (Figure 2). The right arms of the genomes contained a variety of genes including those with DNA replication, processing, and binding functions, membrane proteins, DNA-binding proteins, and numerous genes with no currently known function. The A1 and A5 phages each contained lysin A, lysin B, and holin; the A2 and A6 phages did not have identifiable lysin B genes. Each genome contained an integrase and an immunity repressor, supporting the designation of these phages as temperate.

**Table 2.** Genome characteristics of phages

| Phage Name | Cluster | Genome Length (bp) | GC Content (%) | No. of Protein Coding Genes | No. of tRNAs |
| --- | --- | --- | --- | --- | --- |
| Ajay | A1 | 49,516 | 63.6 | 84 | 0 |
| Wilkins | A1 | 50,907 | 63.9 | 93 | 0 |
| Ph8s | A2 | 52,874 | 62.6 | 98 | 3 |
| AgentM | A5 | 50,503 | 60.9 | 82 | 2 |
| Aragog | A5 | 50,812 | 60.8 | 82 | 1 |
| Archetta | A5 | 47,409 | 60.6 | 76 | 0 |
| ForGetIt | A5 | 51,050 | 60.6 | 85 | 1 |
| Phlorence | A5 | 50,403 | 60.9 | 82 | 1 |
| Koko | A6 | 52,879 | 61.3 | 102 | 3 |

**Figure 2.**
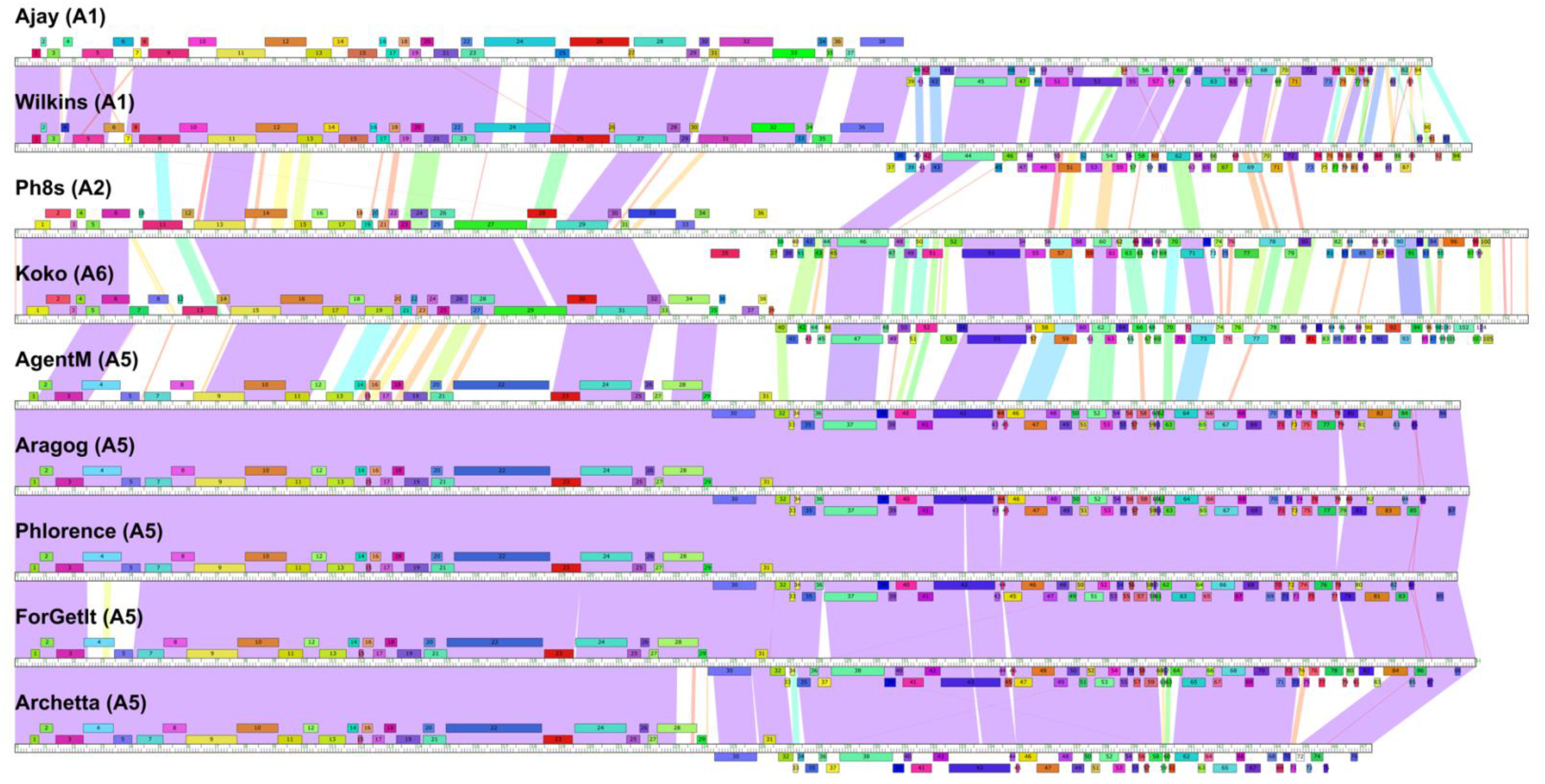
Phamerator map of nine novel Cluster A mycobacteriophages. Subclusters are arranged in order of approximate genomic similarity.

## Discussion

Cluster A, which at the time of publication comprises 662 members, makes up the largest cluster of actinobacteriophages. It has substantial intracluster variation, including unusual and diverse immunity systems [14], and is relatively genetically isolated from other clusters [15,16]. In the present study, nine novel Cluster A mycobacteriophages AgentM, Ajay, Aragog, Archetta, ForGetIt, Koko, Ph8s, Phlorence, and Wilkins were discovered, sequenced, and characterized.

TEM confirmed that these phages, similar to other Cluster A phages, were members of the Siphoviridae family. Genome sequencing and annotation revealed that phages within the same subcluster retained a high degree of genetic similarity, and that all nine phages shared at least some genes with each other. Phages Ph8s and Koko, from subclusters A2 and A6, appear to be more similar to each other than they are to any of the phages from clusters A1 or A5. The isolation and characterization of these phages contributes additional knowledge to the expanding database of actinobacteriophages, and further supports current knowledge of Cluster A phages.

## Acknowledgments

We thank Rebecca A. Garlena and Daniel A. Russell at the Pittsburgh Bacteriophage Institute for phage sequencing and assistance with genome assembly, as well as Debbie Jacobs-Sera, Welkin Pope, Graham Hatfull, and the SEA-PHAGES community for programmatic support. The authors acknowledge the use of instruments at the Electron Imaging Center for NanoMachines supported by NIH (1S10RR23057 to ZHZ) and CNSI at UCLA, as well as the UCLA Genotyping and Sequencing Core, and the NC State Genomic Sciences Laboratory. We would also like to acknowledge the contributions of the following students, teaching assistants and instructors for their roles in isolating, characterizing, and annotating these phages: Samuel Wu, Ryan Ngo, Andrew Lund, Fasih Ahsan, Aida Akopyan, Marianne Arakelyan, Payam Benyamini, Blake Chalman, Alexander Chassiakos, Jasmine Chavez, Michelle Chen, Shirley Cheng, Renee N Crippen, Zain M Dibian, Nam P Do, Oliva Ellis, Hayley Ennis, Megan A FitzPatrick, Chase Fong, Christina Fournier, Kailee Furtado, Joseph Gaballa, Melika Ghalehei, Pirino Giorgia, Emma C Goodwin, Krishna Govindaraju, Ivan Guzman, Mohammad Hamad, Ariel Hamill, Justin Huynh, Joshua Imperial, Rose Jarjoura, Lilith Jiang, Tyrone Johnson, David Joseph, Alyssa Kallman, Christy Kim, Tae W Ko, Antoine Koehl, Sanjan Kumar, Thomas Kwak, Stephen Lai, Gilbert Laim, Yvette M Lakkis, Derek Le, Andrew Lee, Justina Leo, Amber Li, Xiaoying Lian, Rohan Luhar, Catherine Luo, Jenny Luong, Eden Maloney, Josh Martin, Stacee Mendonca, Jordan Meyer, Justin Miller, Samatha Mosier, Ryan Narbutas, Carmen Ng, Christine Nguyen, Teri Nguyen, Daiki Okazaki, Stephanie Orchanian, Jose Ortiz, Emily Pang, Pratik Patel, Cole Pierce, Gloria Y Quach, Viviana Quintana, Vasileios Ragkousis, Krystal Ramos, Ivonne Romero, Cameron Safarpour, Alysha Salbato, Michelle Sanchez, Erin R Sanders, Tasnim Sazzad, Alejandro Serrato-Guillen, Kei T Shao, Bin Shen, Wesley Shen, Alexandra da Silva, Ethan Sim, Rohan Singhal, Emily Smith, Meera Solanki, Jackson Steinberg, Kathleen Tan, Sahar Tashakor, Matthew P Teon, Kevin Tran, Leena Tran, Theresa Tran, Neerja Vashist, Joyce Wang, Larissa Wong, Stephanie Wottrich, Jonathan Yang, Stephanie Yee, Lei Yeh, Edric Yoon, Ashley Yu, Julie Yu, Moses R Zhang, Eric Zhou, Ziqi Zhou, Jovo Vijanderan, William Villella, Giorgia Pirino.

## Author contributions

R.L. and A.F. drafted the paper; A.F and J.M.P. performed quality control on the annotations; A.F. and J.M.P. supervised the research and revised the paper; and all other authors contributed to the isolation, annotation, and genome analysis of the novel phages. This research was funded by the Dean of Life Sciences Division at UCLA, with additional support for sequencing from the HHMI Science Education Alliance-Phage Hunters Advancing Genomics and Evolutionary Science (SEA-PHAGES) program.

## Data availability

The genome sequences of Mycobacterium phages Ajay, AgentM, Aragog, Archetta, ForGetIt, Koko, Ph8s, Phlorence, and Wilkins have been deposited in GenBank under the accession numbers MN062710, MG099934, MG099937, MG099938, MG099944, MG099945, MG099947, MG099949, and MG099951 (NCBI GenBank).

